# Protective effect of *Morus macroura* Miq. fruit extract against acetic acid-induced ulcerative colitis in rats: Involvement of miRNA-223 and TNFα/NFκB/NLRP3 inflammatory pathway

**DOI:** 10.1101/2020.12.22.423927

**Authors:** Rania M. Salama, Samar F. Darwish, Ismail El Shaffei, Noura F. Elmongy, Manal S. Afifi, Ghada A. Abdel-Latif

**Affiliations:** Pharmacology and Toxicology Department, Faculty of Pharmacy, Misr International University (MIU), Cairo, Egypt; Pharmacology and Toxicology Department, Faculty of Pharmacy, Badr University in Cairo, Cairo, Egypt; Biochemistry Department, Faculty of Pharmacy, Misr International University (MIU), Cairo, Egypt; Physiology Department, Faculty of Medicine, Al-Azhar University, Damietta, Egypt; Pharmacognosy Department, Faculty of Pharmacy, Misr International University (MIU), Cairo, Egypt

**Keywords:** Ulcerative colitis, mulberry fruit, TNFα/NFκB, NLRP3, miRNA-223

## Abstract

Tumor necrosis factor receptor (TNFR) activation and nuclear factor kappa B (NFκB) expression play a significant role in the activation of nod-like receptor pyrin domain-1 containing 3 (NLRP3) inflammasome inflammatory pathway, which is involved in the pathogenesis of ulcerative colitis (UC). Furthermore, miRNA-223 expression was shown to exert counter-regulatory effect on NLRP3 expression. Interestingly, polyphenols are attaining increased importance for their potential effectiveness in ameliorating certain diseases owing to their antioxidant and anti-inflammatory activity. In accord, our study attempted to investigate the effect of mulberry tree (*Morus macroura*) fruit extract (MFE) against acetic acid (AA)- induced UC in rats, which is not previously investigated, based on previous promising results for MFE in alleviating gastric ulcer in rats. First, total phenolic (TPC), total flavonoid (TFC), and total anthocyanin content (TAC) were determined in MFE. Then, MFE (300 mg/kg) and sulfasalazine (Sulfa), as a standard treatment (100 mg/kg), were given orally for seven days before intra-rectal induction of UC by AA (2 ml, 4% v/v) on day eight. The extent of UC was evaluated macroscopically and microscopically. Biochemically, the colonic TNFR1, NLRP3, p- NFκB p65, TNFα, interleukin-1 beta (IL-1β), IL-18 levels, miRNA-223 expression and caspase-1 activity were assayed. MFE significantly reduced macroscopic and microscopic scores, colonic levels of TNFR1, NLRP3, p-NFκB p65, TNFα, IL-1β, IL-18, and caspase-1 activity, and showed increased miRNA-223 expression, almost similarly to Sulfa effects. In conclusion, our study provided a novel impact for MFE against AA-induced UC in rats through affecting miRNA-223 expression and halting TNFα/NFκB/NLRP3 inflammatory pathway.

## 1. Introduction

Ulcerative colitis (UC) and Crohn’s disease are the two known categories of inflammatory bowel disease (IBD); a chronic inflammatory disorder that affects the gastrointestinal tract. Though the exact etiology of IBD is not fully clear, however, the immune system is known to be implicated alongside with other gut microbial, genetic and environmental factors, making it a complex multifaceted disorder (Abraham et al., 2017). Unlike Crohn’s disease, UC is characterized by a continuous mucosal inflammation that affects the rectum and extends to the colon as the major sites of injury (Dignass et al., 2012a). UC is considered a problematic disorder as it persists for lifelong, with phases of remission and relapse, in which the patients present with a mix of symptoms such as diarrhea, abdominal pain, rectal bleeding, fatigue and weight loss, thus, impairing their quality of life (Dignass et al., 2012b, Levesque et al., 2015). Moreover, UC increases the risk of colectomy, and on the long-term, colorectal cancer, if not controlled (Van Assche et al., 2013).

One of the major inflammatory pathways involved in the pathogenesis of UC is driven through the key inflammatory mediators; tumor necrosis factor alpha (TNFα) and nuclear factor kappa B (NFκB). TNFα is released in response to harmful stimuli and binds to its receptor (TNFR), which in turn, activates the nuclear factor kappa B (NFκB) heterodimer complex present in the cytosol. This leads to the degradation of the inhibitory IκBα subunit in the NFκB heterodimer complex and the subsequent phosphorylation of the p65 subunit of NFκB and its translocation inside the nucleus (McDaniel et al., 2016, Afonina et al., 2017). P65 is a transcription factor which regulates transcription of multiple genes such as nod-like receptor family pyrin domain-1 containing 3 (NLRP3), pro-interleukin (IL)-1β and pro-IL-18 (Liu et al., 2017). NLRP3 binds to the nascent inflammasome to initiate a downstream signaling cascade via pro-caspase-1, incorporated within this assembled inflammasome complex, which dissociates from the inflammasome complex and becomes activated. Consequently, this active form of caspase-1 converts the inactive substrates pro-IL-1β and pro-IL-18 into the active pro-inflammatory cytokines; IL-1β and IL-18 (Latz et al., 2013). These key inflammatory cytokines significantly trigger the intestinal mucosal inflammation occurring in UC (Zhen and Zhang, 2019). Moreover, NLRP3 inflammasome and caspase-1 effectively contribute to the inflammatory-cell death process, known as pyroptosis (Chen et al., 2019).

Lately, micro RNAs (miRNAs) have gained great attention due to their important role in regulating the expression of various target genes involved in numerous physiological processes. Interestingly, the inflammatory cascade involved in UC can be modulated via miRNA-223, as miRNA-223 was recently reported to inhibit NLRP3 gene expression (Yuan et al., 2018, Kanneganti, 2017). Thus, the development of agents that can inhibit NLRP3 inflammatory pathway, either directly or indirectly through enhancing miRNA-223 expression, can effectively impede the progression of UC.

Moreover, the current remedies of UC show inadequate clinical efficacy and severe side effects (Nielsen and Ainsworth, 2013). Thus, the approach of finding herbal remedies that can treat or halt the progression of mucosal injury and inflammation is gaining much interest in the field of UC. The approach of utilizing herbal remedies comprises plant polyphenols that can be divided into different subgroups; phenolic acids, tannins and flavonoids which include flavonols, flavones, anthocyanins and others (Ozcan et al., 2014). These polyphenolic compounds preserve the ability to act as antioxidants and anti-inflammatory agents, thus, largely contribute to the therapy of multiple inflammatory and auto-immune disorders (Yahfoufi et al., 2018).

Mulberry fruits extract (MFE) is obtained from the completely ripen fruits of mulberry trees (*Morus macroura*), originally native to Pakistan. Multiple beneficial therapeutic effects were attributed to the bioactive ingredients present within MFE, such as flavonoids and phenolic acids (Skrovankova et al., 2015, Nile and Park, 2014). These dietary polyphenolic compounds were accredited for antioxidant and anti-inflammatory activity in peptic ulcer and IBD (Farzaei et al., 2015a, Farzaei et al., 2015b), in addition to their beneficial effects against viruses and bacteria (Daglia, 2012, Cardona et al., 2013). Remarkably, this medicinal plant extract was formerly investigated in gastric ulcer and successfully reduced ulcer indices and inflammation via notable decrease in TNFα (Farrag et al., 2017), however, it was not previously investigated against UC. Therefore, this tempted us to explore for the first time the potential therapeutic benefit of MFE on UC, another form of GIT disorders, through modulating miRNA-223 expression and TNFα/NFκB/NLRP3 signaling cascade which contribute to intestinal inflammation and ulceration.

## 2. Materials and Methods

### 2.1. Ethics statement

The study was conducted in agreement with the ethical procedures and policies approved by the Institutional Review Board of Faculty of Pharmacy, Misr International University, Cairo, Egypt (Approval no.: MIU-PHL2020_03), and conforms with the Guide for the Care and Use of Laboratory Animals (Institute for Laboratory Animal Research, 1996). All efforts were made to minimize animal suffering and to decrease the number of animals used.

### 2.2. Plant material

The fully matured completely ripen *Morus macroura* fruits were freshly collected in April 2020 from Sheikh Zayed City, private garden, Giza, Egypt. The fruits are characterized by a dark purple color and a very sweet taste. They were identified by Prof. Mona M. Marzouk, department of phytochemistry and plant systematics, National Research Centre, Cairo, Egypt. A voucher specimen (M83) had been deposited in the herbarium of National Research Centre (NYBG Steere Herbarium code: CAIRC), Cairo, Egypt. Fresh fruits were dried and kept at 4°C until use.

### 2.3. Animals

Forty adult male Wistar rats (200 ± 20 g) were purchased from The Nile Company for Pharmaceuticals and Chemical Industries (Cairo, Egypt). Rats were allowed one-week acclimatization period at the animal facility of Faculty of Pharmacy; Misr International University in standard polypropylene cages (Four rats per cage). They were allowed free access to normal pellet diet (EL Nasr Pharmaceutical Chemicals Co., Cairo, Egypt) and tap water throughout the experimental period. The rats were kept under standard conditions of temperature (22 ± 2°C) and relative humidity (55 ± 5%) with 12-light/12-dark cycle.

### 2.4. Experimental design

The dose and duration of MFE was adopted from the study of Farrag et al. (2017). MFE was freshly prepared as a solution in distilled water (20 mg/ml). Sulfasalazine (Sulfa) was used as the reference drug, and the dose was obtained from the study of Tekeli et al. (2018). Sulfa was dissolved in normal saline 0.9% to a final concentration of 10 mg/ml.

#### Induction of UC

On day 8 and after overnight fasting, the rats were injected with ketamine (50 mg/kg; i.p.) before intra-rectal administration of 2 ml acetic acid (AA) (Fisher Scientific, Massachusetts, USA), 4% (v/v) in 0.9% NaCl, using a 2.7-mm soft pediatric catheter inserted 8 cm into the colon via the anus (Fatani et al., 2016). Following AA administration, the rats were held vertically for 1 min to prevent AA leakage.

Rats were randomly allocated into 5 groups, each containing 8 rats as follows:

**Group 1:** Rats received single-dose of 2 ml 0.9% NaCl intra-rectally to serve as control.

**Group 2:** Rats received MFE (300 mg/kg/day; p.o.) for 7 days, then single-dose of 2 ml 0.9% NaCl intra-rectally on day 8.

**Group 3:** Rats received single-dose of acetic acid (AA) (2 ml, 4% (v/v) in 0.9% NaCl) intra-rectally to induce UC.

**Group 4:** Rats received MFE (300 mg/kg/day; p.o.) for 7 days, then single-dose of AA (2 ml, 4% (v/v) in 0.9% NaCl) intra-rectally on day 8.

**Group 5:** Rats received Sulfa (100 mg/kg/day; p.o.) for 7 days, then single-dose of AA (2 ml of 4% (v/v) in 0.9% NaCl) intra-rectally on day 8.

#### Tissue collection

On day nine, all animals were anesthetized with a cocktail of i.p. ketamine hydrochloride (50 mg/kg) and xylazine (5 mg/kg) (Koc et al., 2005) purchased from Sigma-Aldrich (Missouri, USA), then sacrificed and their colon tissues were quickly dissected, examined macroscopically and the number and area of the ulcers were recorded. Subsequently, each colon was cut into small pieces; one piece (5 cm long) was cut from each colon and rapidly flushed and fixed in 10% neutral buffered formalin for 72 hours and processed for light microscopical examination. The other pieces of the colon segment from each rat were divided into portions and immediately stored at −80°C until assessment of biochemical markers. All assessments were carried out by blinded investigators.

### 2.5. Methods

#### 2.5.1. Preparation of the mulberry fruit extract

Two hundred grams of the fruits of *Morus macroura* were extracted for three times by 70% ethanol 1:10 (w/v) in an ultrasonic bath for 45 min at 50°C. The extracts were filtered through Whatman^®^ Grade 1 filter paper, combined together, and then dried using a rotary evaporator under vacuum at 40°C. The dried MFE was lyophilized and stored in a refrigerator at 4°C until required for use. For biological studies, the extract was freshly prepared as a suspension in distilled water before use.

#### 2.5.2. Determination of polyphenolic compounds

Since polyphenols are among the most biologically active constituents, total phenolic content (TPC), total flavonoid content (TFC), and total anthocyanin content (TAC) were determined in the MFE. TPC was determined by colorimetric assay, using the Folin–Ciocalteu colorimetric method (Singleton and Rossi, 1965, Singleton et al., 1999). The TPC was expressed as mg gallic acid equivalents (GAE)/g extract.

TFC was quantified by colorimetric assay, using the method described by Chang et al. (2002). The TFC was expressed as mg rutin equivalents/g extract.

TAC was determined by the pH-differential method (Giusti and Wrolstad, 2001). Anthocyanin pigments undergo reversible structural transformations with a change in pH manifested by strikingly different absorbance spectra. TAC was expressed as milligrams cyanidin-3-glucoside equivalents per gram of dry weight purification (mg C-3-G/g DW).

#### 2.5.3. Evaluation of disease activity index (DAI)

The disease activity index (DAI) is calculated based on the cumulative scores of percent loss in body weight (0-4), stool consistency (0-4) and gross bleeding (0-4), and then divided by 3 as previously described by Cooper et al. (1993).

Body weight loss was calculated as the percentage difference between body weight before UC induction (Day 8) and the final body weight before sacrifice (Day 9). For stool consistency, it was scored as follows: (0: normal, 1 and 2: loose stool, 3 and 4: diarrhea). The presence of occult blood in feces was assayed using benzidine test (Kiefer, 1934). Occult blood was graded using a score of 0, for no color change; 1, for a very light blue color (±) taking over 30 s to appear; 2, for a blue color developing in 30 s or more (+); 3, for an immediate change in color occurring in less than 30 s (++); and 4, for gross blood observable on the slide (Table 1).

**Table 1.** Disease activity index (DAI) score evaluation of ulcerative colitis (UC) in rats.

| DAI score | % weight loss | Stool consistency | Rectal bleeding |
| --- | --- | --- | --- |
| 0 | None | Normal | Negative |
| 1 | 1-5% | | Occult blood ± |
| 2 | 6-10% | Loose stool | Occult blood + |
| 3 | 11-20% |  | Occult blood ++ |
| 4 | > 20% | Diarrhea | Gross blood |

#### 2.5.4. Macroscopic assessment of colitis

The UC induced in the experimental animals was evaluated based on its macroscopic characteristics as determined by the previously described scoring system by Saber et al. (2019). The macroscopic damage was scored by a blinded investigator according to the following scale: 0 = no macroscopic change, 1 = mucosal erythema only, 2 = mild mucosal edema, slight bleeding, or small erosions, 3 = moderate edema, slight bleeding ulcers or erosions and 4 = severe ulceration, erosions, edema, and tissue necrosis.

#### 2.5.5. Evaluation of ulcer area (UA) and ulcer index (UI)

Ulcer area (UA) was measured using plain glass square. Each cell on the glass square was 1mm^2^ in area, and the number of cells was counted, and the ulcer area was determined for each colon.

Ulcer index (UI) was determined as described earlier (Ozbakis Dengiz and Gursan, 2005) using the following equation;

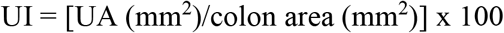

#### 2.5.6. Assessment of colonic inflammatory markers

Rat sandwich ELISA kit was purchased from MyBioSource (California, USA) for the assay of IL-1β (catalogue # MBS825017) and IL-18 (catalogue # MBS355269). Similarly, rat ELISA kit was purchased from Cusabio (Wuhan, P.R. China) to assess TNFα (catalogue # CSB-E11987r). All procedures were done according to the manufacturers’ instructions.

#### 2.5.7. Assessment of miRNA-223 expression in colon tissue

Total miRNA was extracted from colon tissue homogenate using mirVana™ miRNA Isolation Kit (Ambion®, Texas, USA). The extracted miRNA was reverse transcribed to cDNA using miScript II RT Kit (Qiagen, Hilden, Germany) according to the manufacturer’s instructions. Quantification of the target miRNA-223 “GenBank accession no.: **NR_031936.1**” from cDNA was performed by real-time PCR using the miScript SYBR® Green PCR Kit (Qiagen, Hilden, Germany) as described by the manufacturer (*Forward primer*: 5’ TGGCCTTCTGCAGTGTTACG 3’-*Reverse primer*: 5’ AAGCATGAGCCACACTTGGG 3’). The relative expression of miRNA was obtained using the ∆∆ CT method as previously described (Livak and Schmittgen, 2001) using U6 “GenBank accession no.: **K00784.1**” as a housekeeping gene (*Forward primer*: 5’ CTCGCTTCGGCAGCACA 3’-*Reverse primer*: 5’ AACGCTTCACGAATTTGCGT 3’).

#### 2.5.8. Western blot analysis

Samples of equal protein concentrations (≈ 20 μg) were electrophoresed using 10% Sodium Dodecyl Sulfate/polyacrylamide gel (SDS/PAGE) and electro-transferred to polyvinylidene difluoride membranes. The membranes were blocked with 5% (w/v) skimmed milk powder in PBS/Tween-20 for 2 hours at room temperature. Then, the membranes were incubated with TNFR1 polyclonal 1:1000 (Abcam, catalogue # ab19139), p-NFκB p65 (S536) polyclonal 1:2000 (Abcam, catalogue # ab86299), total NFκB p65 polyclonal 1:2000 (Abcam, catalogue # ab16502), NLRP3 polyclonal 1:1000 (Abcam, catalogue # ab214185), pro- and cleaved caspase-1 monoclonal 1:1000 (Abcam, catalogue # ab179515) antibodies diluted in tris-buffered saline-tween containing 1% bovine serum albumin and β-actin (Santa Cruz Biotechnology) as internal control diluted 1:1000 in blocking buffer. The membranes were incubated with the corresponding secondary antibodies for 1 hour at room temperature, washed, and then developed. Finally, images of indicated protein bands were recorded on the BioMax film (Kodak), and densitometrical quantification was conducted by using Image J software (Bio-Rad, California, USA). For quantitative purposes, the OD values of phospho-NFκB p65 was normalized to the detection of non-phospho NFκB p65 in the same sample, and cleaved caspase-1 was normalized to the detection of pro-caspase-1 in the same sample. Densities of TNFR1 and NLRP3 bands were standardized to the corresponding density of β-actin. Data were expressed as a fold change of the control group.

#### 2.5.9. Protein content

Protein concentration in colon tissue samples was estimated as described by Bradford (1976).

#### 2.5.10. Light microscopic examination

After fixation of the colon, the tissues were washed in several changes of 70% ethanol, followed by dehydration in ascending grades of alcohol, clearing in xylene and embedding in paraffin wax to obtain paraffin blocks (Suvarna et al., 2018). Colonic sections of 5μm thickness were cut, mounted on slides, stained with Hematoxylin and Eosin (H&E) and examined by histopathologist, blinded to all treatments. Severity of lesions was graded on a scale of negative (-) to severe (+++).

### 2.6. Statistical Analysis

Each variable was tested for normality using the Kolmogorov-Smirnov test. All data were normally distributed, thus, followed parametric analysis and were expressed as means ± SD and compared using one-way ANOVA followed by Tukey’s *post hoc* test. Level of probability (*p* value) less than 0.05 is used as the criterion of significance. Statistical analysis was performed using the statistical software package GraphPad Prism^®^, Version 5.00 for Windows (California, USA).

## 3. Results

The statistical comparison between control and MFE (300 mg/kg/day), revealed no significant difference; therefore, all comparisons were referred to the control group.

### 3.1. Assessment of TPC, TFC, and TAC of MFE

The results of the determination of TPC, TFC, and TAC in the studied extract (MFE) are presented in Table 2, which shows that the levels of each of them are high.

**Table 2.** Total phenolic content (TPC), total flavonoid content (TFC), and total anthocyanin content (TAC) of *Morus macroura* fruit extract (MFE)

| Sample Code | % extraction<br>yield | TPC <sup>a</sup> (mg/g) | TFC <sup>b</sup> (mg/g) | TAC <sup>c</sup> (mg/g) |
| --- | --- | --- | --- | --- |
| MFE | 15.3±1.2 | 51.03±2.81 | 34.16±1.94 | 4.21± 0.44 |

Values (mean ± SD) are averages of three samples of each fruit, analyzed individually in triplicate ( $p < 0.05$ ); <sup>a</sup> as per gallic acid equivalent (mg GAE/g DW); <sup>b</sup> as per rutin equivalent (mg rutin /g DW) and <sup>c</sup> as per as cyanidin-3-glucoside equivalent (mg C-3-G/g DW).

### 3.2. Evaluation of MFE effect on DAI

As shown in Fig.1A, overt rectal bleeding, one of the scoring parameters of DAI, was observed in AA-treated rats only. Upon calculating DAI [F (4, 32) = 28.16, *p* ˂ 0.001], significant increase in the DAI was observed in the AA-treated group by 19-fold, when compared to control rats (Table 3). This index was markedly decreased in groups pre-treated with MFE and Sulfa by 47.9% and 36.8%, respectively, when compared to AA-treated group. No significant difference was observed between MFE- and Sulfa-treated groups.

**Fig. 1.**
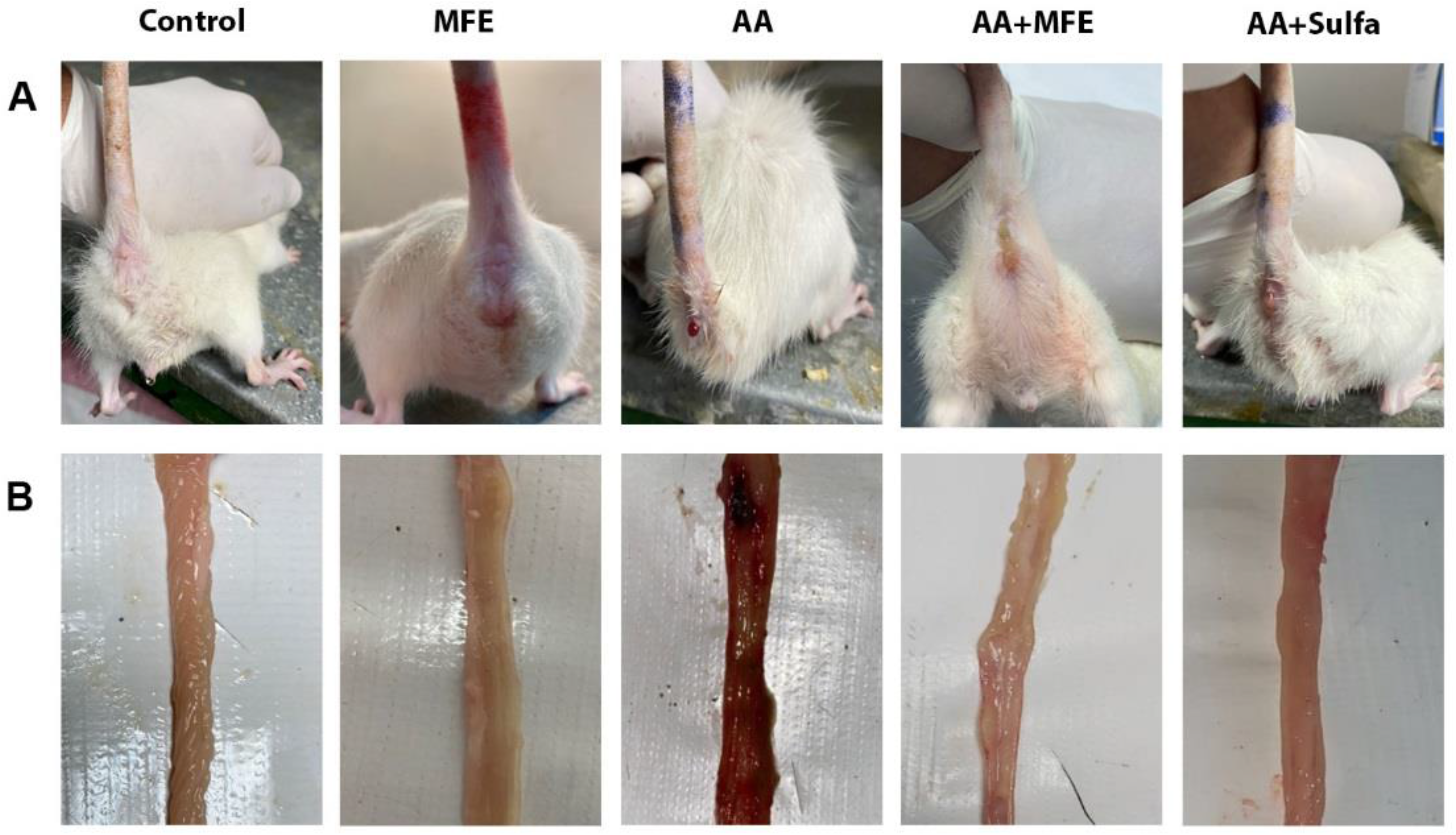
(A) Representative photographs of rats’ anal region for observation of rectal bleeding, and (B) the dissected colons for macroscopic evaluation of colon damage in normal control, MFE-treated group (300 mg/kg/day; p.o.), AA-treated group for induction of UC (2 ml, 4% (v/v) in 0.9% NaCl; intra-rectally), AA+MFE-treated group, and AA+Sulfa-treated standard group (100 mg/kg/day; p.o.). AA, acetic acid; MFE, mulberry fruit extract; Sulfa, sulfasalazine; UC, ulcerative colitis.

**Table 3.** Effect of MFE on DAI, macroscopic score, ulcer area and ulcer index in AA-induced UC in rats.

| Groups | DAI | Macroscopic score | Ulcer area (mm<sup>2</sup>) | Ulcer index |
| --- | --- | --- | --- | --- |
| Control | 0.125 ± 0.17 | 0.0 ± 0.0 | 0.0 ± 0.0 | 0.0 ± 0.0 |
| MFE | 0.0 ± 0.0 | 0.0 ± 0.0 | 0.0 ± 0.0 | 0.0 ± 0.0 |
| AA | 2.375 ± 0.58*** | 3.375 ± 0.52*** | 523.8 ± 185*** | 60.52 ± 16.2*** |
| AA+MFE | 1.238 ± 0.85## | 2.286 ± 0.76### | 204.7 ± 218.7## | 15.41 ± 15.32### |
| AA+Sulfa | 1.5 ± 0.62# | 2.333 ± 0.52## | 210.8 ± 227.4## | 22.10 ± 29.10### |

Data are presented as the mean ± SD (n = 8 per group; one-way ANOVA followed by Tukey's multiple comparison test; \*\*\* $p < 0.001$ , vs. the control group; # $p < 0.05$ , ## $p < 0.01$ , ### $p < 0.001$ vs. the AA-treated group). AA, acetic acid; DAI, disease activity index; MFE, mulberry fruit extract; Sulfa, sulfasalazine; UC, ulcerative colitis.

### 3.3. Macroscopic evaluation of colonic tissues following MFE treatment

As shown in table 3 and Fig. 1B, the macroscopic examination score [F (4, 32) = 87.45, *p* ˂ 0.001] of the dissected colon tissue was significantly higher in AA-treated rats, when compared to the control rats. This score was markedly decreased in both MFE and Sulfa pre-treated groups by 32.3% and 30.9%, respectively, when compared to AA-treated group. However, no significant difference was shown between MFE- and Sulfa-treated groups.

### 3.4. Effect of MFE on UA and UI

The UA [F (4, 32) = 15, *p* < 0.001] and UI [F (4, 32) = 21.07, *p* < 0.001] were significantly increased upon AA treatment, when compared to control rats. The UA was markedly decreased in MFE and Sulfa pre-treated rats by 60.9% and 59.8%, respectively, when compared to AA-treated group. Similarly, the UI was significantly reduced following MFE and Sulfa pre-treatment by 74.5% and 63.5%, respectively, when compared to its value in AA-treated rats (Table 3). Nevertheless, there was no significant difference between MFE and Sulfa pre-treatment regarding their effect on both UA and UI.

### 3.5. Effect of MFE on miRNA-223 gene expression and inflammatory markers levels in colon tissues

As shown in Fig. 2A, relative expression of miRNA-223 [F (4, 35) = 66.62, *p* < 0.001] was significantly reduced upon AA treatment by 65.3%, when compared to the expression in the control group. Inversely, the inflammatory markers TNFα [F (4, 35) = 210.2, *p* < 0.001], IL-1β [F (4, 35) = 141.6, *p* < 0.001], and IL-18 [F (4, 35) = 194.3, *p* < 0.001] levels were significantly increased upon AA treatment by 5.3, 4.4 and 3 folds, respectively, when compared to those in control rats (Fig. 2B-2D). Such effects were improved by MFE pre-treatment, where 2.3-fold significant increase in miRNA-223 expression was shown, while the inflammatory markers levels were significantly reduced as follows; TNFα (64.8%), IL-1β (50%), and IL-18 (54.8%), when compared to the AA-treated group. Similarly, pre-treatment with the standard drug Sulfa showed 2.1-fold significant increase in miRNA-223 expression, with significant reduction in the inflammatory markers as follows; TNFα (59.3%), IL-1β (48.5%), and IL-18 (55.3%), when compared to the AA-treated group. Yet, no significant difference was observed in these markers between MFE- and Sulfa-treated groups.

**Fig. 2.**
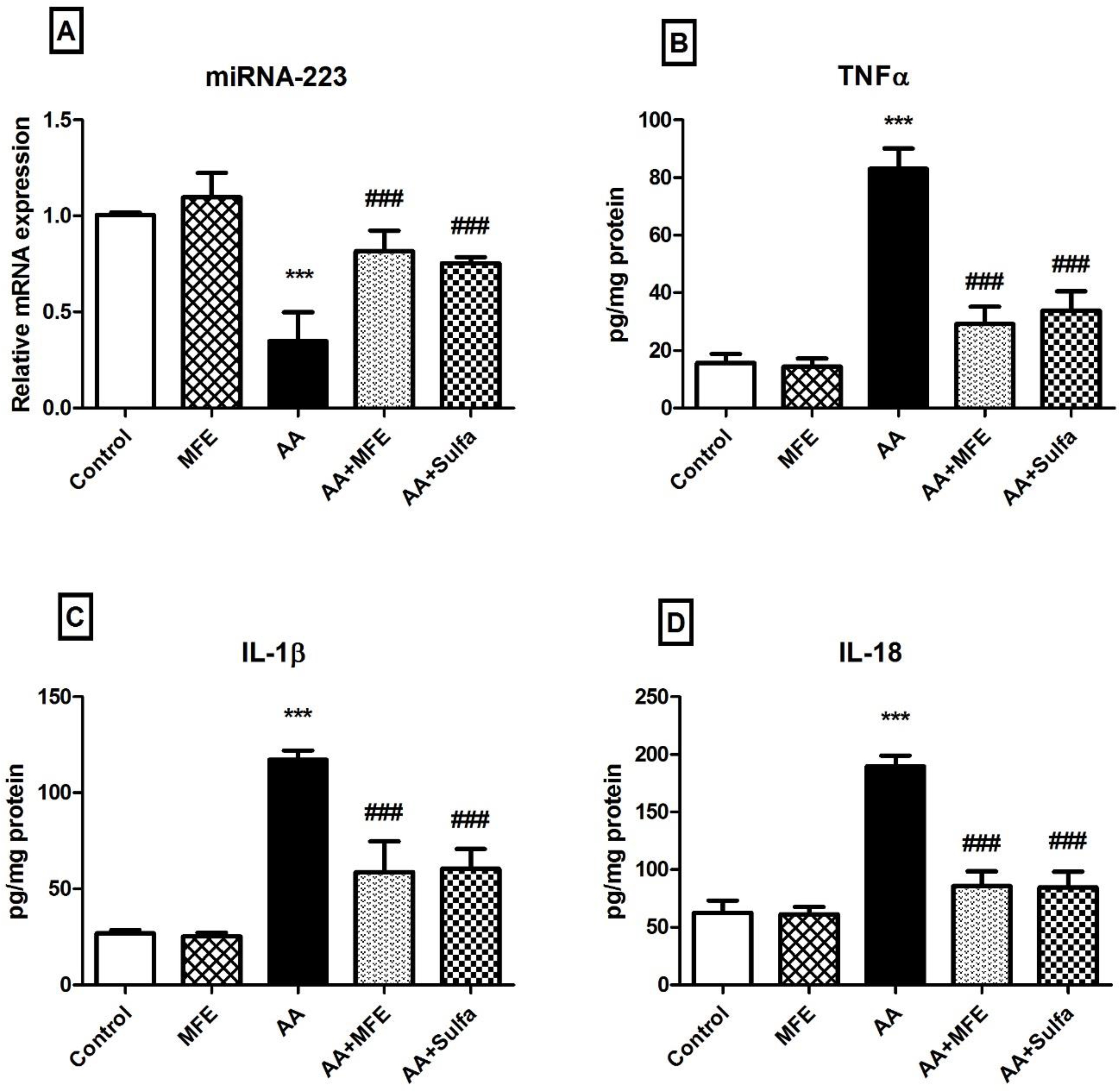
Effect of MFE on (A) miRNA-223 gene expression and protein levels of (B) TNFα, (C) IL-1β and (D) IL-18 in rat colons. Data are presented as the mean ± SD (n = 8 per group; one-way ANOVA followed by Tukey’s multiple comparison test; *** *p* < 0.001, vs. the control group; ^###^ *p* < 0.001, vs. the AA-treated group). AA, acetic acid; IL, interleukin; MFE, mulberry fruit extract; Sulfa, sulfasalazine; TNFα, tumor necrosis factor alpha.

### 3.6. Effect of MFE on protein expression of TNFR1, p-NFκB p65 (S536), NLRP3 and caspase-1 in colon tissues

The levels of TNFR1 [F (4, 35) = 59.05, *p* < 0.001], p-NFκB p65 (S536)/total NFκB p65 [F (4, 35) = 297.3, *p* < 0.001], NLRP3 [F (4, 35) = 283, *p* < 0.001] and cleaved caspase-1/caspase-1 [F (4, 35) = 143.8, *p* < 0.001] were significantly increased in the AA-treated group by 5.6, 5.3, 6 and 4.9 folds, respectively, when compared to those in control rats (Fig. 3). Following treatment with MFE, TNFR1, p-NFκB p65 (S536), NLRP3 and caspase-1 relative protein expression levels were significantly reduced by 62.1%, 63.1 %, 57.7% and 61.6%, respectively, when compared to the AA-treated group. Likewise, pre-treatment with Sulfa showed significant reduction in TNFR1, p-NFκB p65 (S536), NLRP3 and caspase-1 relative protein expression levels by 56.4%, 53.2 %, 52.4% and 59.5%, respectively, when compared to the AA-treated group. Notably, MFE showed significantly lower p-NFκB p65 (S536) when compared to Sulfa-treated group by 21.3%. However, no significant difference was observed between MFE- and Sulfa-treated groups in TNFR1, NLRP3 and caspase-1 protein expression levels.

**Fig. 3.**
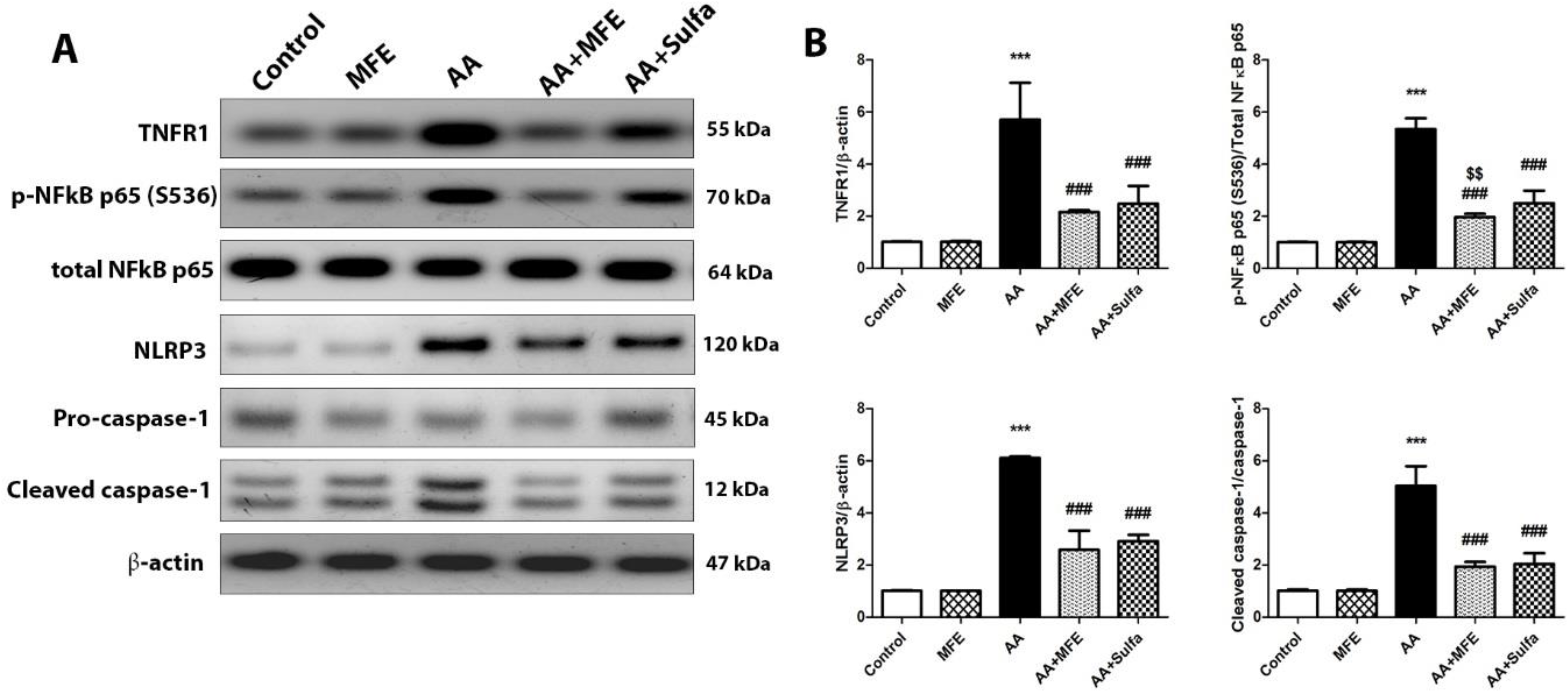
(A) Representative western blot bands of protein levels. (B) Quantitation of TNFR1, phospho-NFκB p65 (S536)/total NFκB p65, NLRP3 and cleaved caspase-1/caspase-1 in rat colons. Data are presented as the mean ± SD (n = 8 per group; one-way ANOVA followed by Tukey’s multiple comparison test; ^***^ *p* < 0.001, vs. the control group; ^###^ *p* < 0.001, vs. the AA-treated group; ^$$^ *p* < 0.01 vs. Sulfa group). AA, acetic acid; MFE, mulberry fruit extract; NFκB, nuclear factor kappa B; NLRP3, nod-like receptor pyrin domain-1 containing 3; Sulfa, sulfasalazine; TNFR, tumor necrosis factor receptor.

### 3.7. Histopathological results of light microscopic examination of the colon sections

As shown in Fig. 4 and table 4, microscopic examination of colon tissue sections of the control group demonstrated normal histological structures of intestinal wall with intact lining mucosa including intact intestinal crypts with normal densities of goblet cells, intact submucosal layer as well as outer muscular coat. Similarly, colon sections from rats treated with MFE only showed intact intestinal wall without abnormal structural changes.

**Table 4.** Histopathologic microscopic examination of the rat colons.

|  | Control | MFE | AA | AA+MFE | AA+Sulfa |
| --- | --- | --- | --- | --- | --- |
| **Erosions/ulcerations** | - | - | +++ | + | + |
| **Inflammatory cells infiltrates** | - | + | +++ | ++ | ++ |
| **Edema** | - | - | ++ | + | + |
| **Hemorrhage** | - | - | +++ | - | - |
| **Outer muscular coat necrotic changes** | - | - | +++ | - | - |

- = negative; + = mild (less than 20% of samples); ++ = moderate (21-50% of samples); +++ = severe (more than 50% of samples). AA, acetic acid; MFE, mulberry fruit extract; Sulfa, sulfasalazine.

**Fig. 4.**
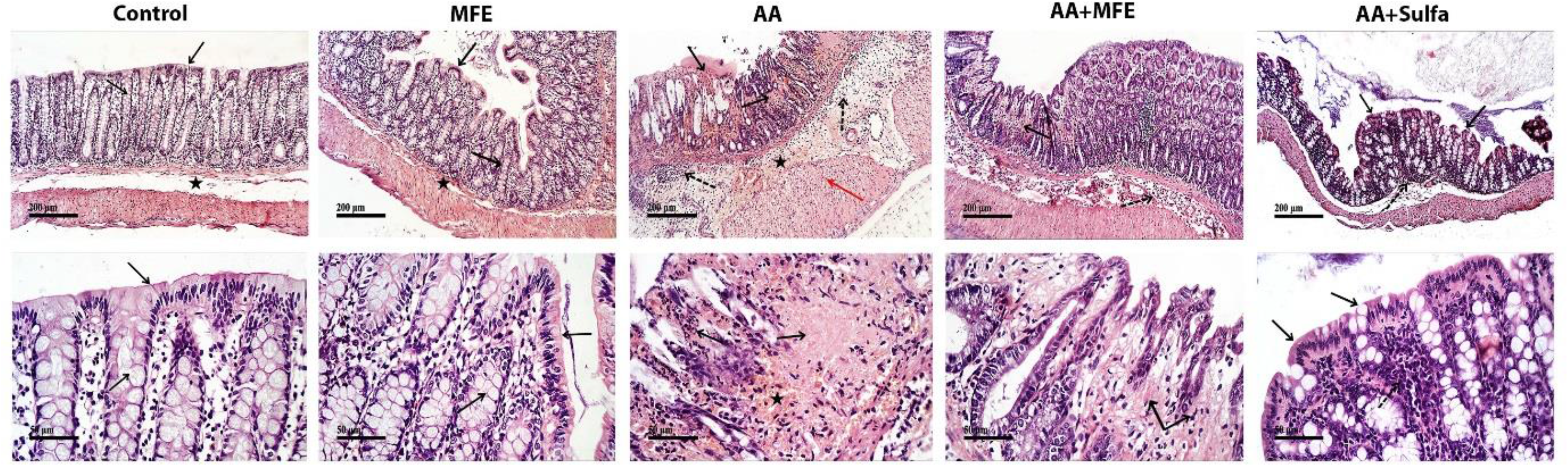
Photomicrographs of colon sections from control group showing normal histological structures with normal densities of goblet cells (black arrow), intact submucosal layer and outer muscular coat (star). Mulberry fruit extract (MFE) group showed intact intestinal wall without abnormal structural changes, similar to control group. Acetic acid (AA) group showed severe mucosal hemorrhagic ulcerative lesions with abundant necrotic tissues (black arrow), submucosal inflammatory cells infiltrates (dashed arrow), hemorrhagic patches (star), and severe necrotic changes of outer muscular coat (red arrow). Pre-treatment with MFE (AA+MFE group) demonstrated significant reduction of mucosal ulceration with occasional records of focal areas of necrotic tissues (black arrow) and mild decrease of inflammatory cells infiltrates (dashed arrow). Pre-treatment with sulfasalazine (Sulfa) (AA+Sulfa group) showed almost intact lining mucosa (black arrow) with moderate mucosal and submucosal inflammatory cells infiltrates (dashed arrow). (H&E stain, x100, x400)

Inversely, colon sections of AA-treated rats showed persistence of wide severe mucosal hemorrhagic ulcerative lesions with abundant necrotic tissue depress records as well as submucosal inflammatory cells infiltrates, hemorrhagic patches and edema, in addition to severe necrotic changes of outer muscular coat.

Interestingly, pre-treatment of rats with MFE demonstrated significant reduction of mucosal ulceration records with occasional records of focal areas of necrotic tissue depress as well as diminished records of mature goblet cells. Moderate submucosal edema was evident accompanied with mild decrease of inflammatory cells infiltrates. Intact outer muscular coat was also observed.

Nearly similar picture was observed in rats pre-treated with Sulfa, in which colon sections showed almost intact lining mucosa with mild focal scattered epithelial erosions. Mild records of submucosal edema with moderate mucosal and submucosal inflammatory cells infiltrates and intact outer muscular coat were also noted.

## 4. Discussion

The pathogenesis of UC has been disclosed gradually. However, the effective way to prevent and treat this inflammatory disease remains elusive (Wang et al., 2019) and the long term treatment shows limited effectiveness and failure of response, in addition to serious side effects. Accordingly, the development of new treatment regimens and the exploration of new natural products with minimal side effects, for alleviation of this inflammatory disease is crucial (Lean et al., 2015).

On this basis, we were inspired to explore for the first time the potential therapeutic benefit of the MFE on UC as it was previously shown to be rich in polyphenolic compounds possessing anti-inflammatory and antioxidant effects (Farrag et al., 2017). Flavonoids and anthocyanins belong to the phenolic compounds class, and in this study high level of phenolic compounds produced high levels of flavonoids and anthocyanins, which indicates that they are the major contributors to the presence of phenolics in MFE. In addition, the results of the determination of polyphenolic contents showed that the TPC of MFE was significantly higher than the results performed in Pakistan (Mahmood et al., 2012) and in Ismailia, Egypt (Khalil and Embaby, 2017). This may be due to the effect of the differences in ripening stages, as well as the environmental conditions and soil in which the plant was grown. Additionally, this may be due to the ultrasonic treatment that was used for extraction, as it showed a significant increase in extraction efficiency for preparing an antioxidant-rich fruit extract from mulberry fruit. Moreover, it showed some advantages, such as shorter extraction time and higher extraction yield for TPC and TAC compared to normal processing (Nguyen and Nguyen Ha, 2018).

Herein, the protective effect of MFE against AA-induced UC in rats was investigated through measuring the disease activity markers such as the DAI, the UA and UI, in addition to the colonic macroscopic and microscopic examination, together with exploring the inflammatory signaling pathway and the molecular targets of MFE. Sulfasalazine was used as a reference drug for the protective effect against UC.

Our results confirmed that the pre-treatment of rats with MFE before administration of AA had a protective effect against UC, which was evidenced by the marked reduction in the DAI, the UA and the UI. In addition, reduction in the macroscopic examination score, preservation of the intestinal mucosa and apparent reduction in the colon tissue inflammation was also observed. Moreover, the inflammatory signaling cascade was inhibited through the downregulation of TNFR1 and the overexpression of miRNA-223 which led to the subsequent inhibition of the TNFα/p-NFκB p65/NLRP3/caspase-1 signaling pathway. Hence, our study stated for the first time the protective effect of MFE against UC in rats.

While comparing the AA-induced UC rats with the control ones, there was a marked increase in the DAI, UA, UI and the macroscopic examination score. Furthermore, the histopathological examination revealed an apparent increase in the mucosal ulceration, hemorrhage, and necrosis in addition to submucosal inflammatory cells infiltration, edema and necrotic lesions in the outer muscular layer. These findings were consistent with the results of previous studies (El-Far et al., 2020, Owusu et al., 2020). Interestingly, the pre-treatment of rats with MFE showed a significant reduction in the DAI, UA and the UI compared to the untreated AA-induced UC rats. Moreover, the histopathological examination confirmed the preservation of the colonic integrity due to the apparent reduction in the mucosal ulceration, edema and necrotic changes together with the significant reduction in the inflammatory cells infiltration. The effect of MFE on UC had not been explored before, however, it was tested in a previous study against gastric ulcer in rats and showed similar findings, where the protective anti-ulcer effect was confirmed following pre-treatment with MFE due to its antioxidant and anti-inflammatory effects (Farrag et al., 2017). Another study was performed using the mulberry big leaf extract of the same plant; *Morus macroura* and also proved its gastroprotective effect in a model of water immersion and restraint stress-induced gastric ulcer in mice (Wei et al., 2018). Furthermore, several studies were performed using the MFE prepared from other species such as *Morus alba* and *Morus nigra*, which are also rich in polyphenolic compounds and the results proved the gastroprotective effect of these extracts due to its anti-inflammatory effect (Chandra et al., 2016, Nesello et al., 2017). Taken together all the consistent previous findings with the current results, it was suggested that the protective effect of MFE against UC is most probably due to the presence of high content of phenolic compounds, which might have possessed a strong anti-inflammatory effect.

In an attempt to reveal the possible mechanism of the anti-inflammatory effect of MFE, the current study focused on the investigation of TNFα and NFκB signaling cascades, because these markers are among the key players of the inflammatory response involved in the pathogenesis of UC, playing a major role in the regulation of expression of the pro-inflammatory cytokines such as IL-1β, and IL-6 (Wang et al., 2017, El-Far et al., 2020). Additionally, a recent review highlighted the role of TNFα and TNFR1 activation in the progression of the inflammatory response and consequent activation of NFκB signaling pathway in different inflammatory diseases including UC (Holbrook et al., 2019). It was also reported in numerous literatures that NFκB pathway plays a central role in regulating the release of cytokines and contributes to the inflammatory and immune response in patients with UC (Sakthivel and Guruvayoorappan, 2014, Wang et al., 2015), which was consistent with our findings where the induction of UC in rats using AA increased the level TNFα, TNFR1 that consequently augmented the activation of NFκB p65.

Intriguingly, the pre-treatment with MFE led to reduction in the levels of TNFα and TNFR1, causing a significant reduction in the level of the p-NFκB p65. Similar results were previously reported where the anti-inflammatory effect of sanguinarine protected the colon against AA-induced UC in rats through inhibition of TNFα and consequent inhibition of p-NFκB p65 (Niu et al., 2013). Another study provided evidence for the promising therapeutic effect of galangin in cyclical DSS‐induced colitis through the attenuation of p-NFκB p65 activation and lowering the levels of TNFα (Gerges et al., 2020). Furthermore, a recent review summarized the Chinese medicines used in the treatment of UC and provided numerous evidences on the role of the attenuation of TNFα and p-NFκB p65 in UC treatment (Lu and Zhao, 2020). These results, collectively with the current findings, led us to suggest that the anti-inflammatory effect of MFE was probably executed through the inhibition of the TNFα/p-NFκB p65 signaling pathway.

Activation of p-NFκB p65 is considered the first step in the activation of the NLRP3 inflammasome which was proved to be responsible for the activation of caspase-1. In turn, caspase-1 activates IL-1β and IL-18, which stimulate further release of other inflammatory cytokines such as IL-6 and TNF α (Saber and El-Kader, 2020, Yang et al., 2019). This cascade was confirmed by previous work where the NLRP3 inflammasome was activated in response to the activation of NFκB and followed by the activation of caspase-1, IL-1β and IL-18 in a model of DSS-induced UC in rats (Saber and El-Kader, 2020). Another study showed the same results, both in vivo using a model of DSS-induced UC in mice, and in vitro using LPS-induced RAW264.7 cells (Qiao et al., 2020). Our results were consistent with the aforementioned findings though using a different method of UC induction, and to date this pathway was not explored in our model of AA-induced UC. These data again led us to postulate that targeting this inflammatory cascade could have a beneficial effect in the treatment of UC.

Impressively, pre-treatment with MFE caused an apparent decrease in the levels of NLRP3, caspase-1, IL-1β and IL-18 which is probably due to the inhibitory effect of MFE on the TNFα/p-NFκB p65 pathway. Similarly, Qiao et al. showed that JWSYD effectively treated DSS-induced UC by inhibiting the NLRP3 inflammasome and NFκB pathway (Qiao et al., 2020). Another study also reported the beneficial effects of xanthine oxidase inhibitors in the treatment of DSS-induced UC in rats via inhibition of IL-1β and IL-18 (El-Mahdy et al., 2020). Moreover, Cheng et al. stated that isoorientin; a C-glycosylflavone showed a promising effect in the treatment of trinitrobenzene sulfonic acid-induced bowel disease through the amelioration of NLRP3 (Cheng et al., 2020). Hence, by referring to these previous results together with the current data we suggested that MFE exhibited its beneficial anti-inflammatory effect via the NFκB dependent inhibition of NLRP3 inflammasome which in turn stopped the inflammatory response by inhibition of the cleavage of caspase-1, pro-IL-1β and pro-IL-18.

Recent studies reported the downregulation of miRNA-223 in IBD (Din et al., 2020, Konstantinidis et al., 2020, Wu et al., 2020). This was consistent with our findings, in which miRNA-223 expression was significantly decreased following AA-induction of UC in rats. Several studies conducted on different types of inflammatory disease confirmed that the upregulation of miRNA-223 resulted in the inhibition of NLRP3 and consequent inhibition of the inflammatory cytokines IL-1β and IL-18 (Din et al., 2020, Jimenez Calvente et al., 2020, Wu et al., 2020, Zhang et al., 2020), which encouraged us for the first time to investigate the impact of miRNA-223 on the inflammatory response in AA-induced UC. Herein, the pre-treatment with MFE markedly increased the relative expression of miRNA-223, resulting in a marked reduction in the NLRP3 in addition to the inhibition of caspase-1, IL-1β and IL-18 as discussed above. However, Kim et al. showed that miRNA-223 was upregulated in IBD and that this upregulation resulted in the activation of NFκB (Kim et al., 2016). On the other hand, recent studies supported our results, where Wu et al. proved that the silencing of miRNA-223 resulted in the upregulation of NLRP3, while overexpression of miRNA-223 resulted in the inhibition of NLRP3 and the consequent inhibition of caspase-1, IL-1β, IL-18 and pyroptosis inhibition (Wu et al., 2020). Additionally, Din et al. provided evidence that Bifidobacterium bifidum possessed a strong anti-inflammatory effect against DSS-induced UC in mice through the upregulation of miRNA-223, which suppressed the level of NLRP3 (Din et al., 2020). These results collectively with our data led us to propose that the MFE might have exerted its anti-inflammatory effect against UC through the upregulation of miRNA-223 which contributed to the inhibition of the NLRP3 inflammasome.

The anti-inflammatory effect of MFE against AA-induced UC in rats was compared to Sulfa as a reference drug and all the findings showed no significant difference between both treatments, except for the level of p-NFκB p65, where MFE had a stronger inhibitory effect on the p-NFκB p65 level than Sulfa. This was probably due to the dual mechanism of the MFE as we suggested before, that it inhibited the p-NFκB p65 via inhibition of TNFα, TNFR1 on one hand, and on the other hand by the downregulation of miRNA-223 which was previously reported to have a negative regulatory effect on the NFκB signaling pathway (Roffel et al., 2020).

In conclusion, to our knowledge the results of the current study shed the light for the first time on the beneficial therapeutic effect of MFE against AA-induced UC through the inhibition of the inflammatory response by a mechanism that might involve a crosstalk between the miRNA-223, NFκB and the NLRP3 inflammasome. Further studies are needed to provide more evidence on the suggested mechanism herein.

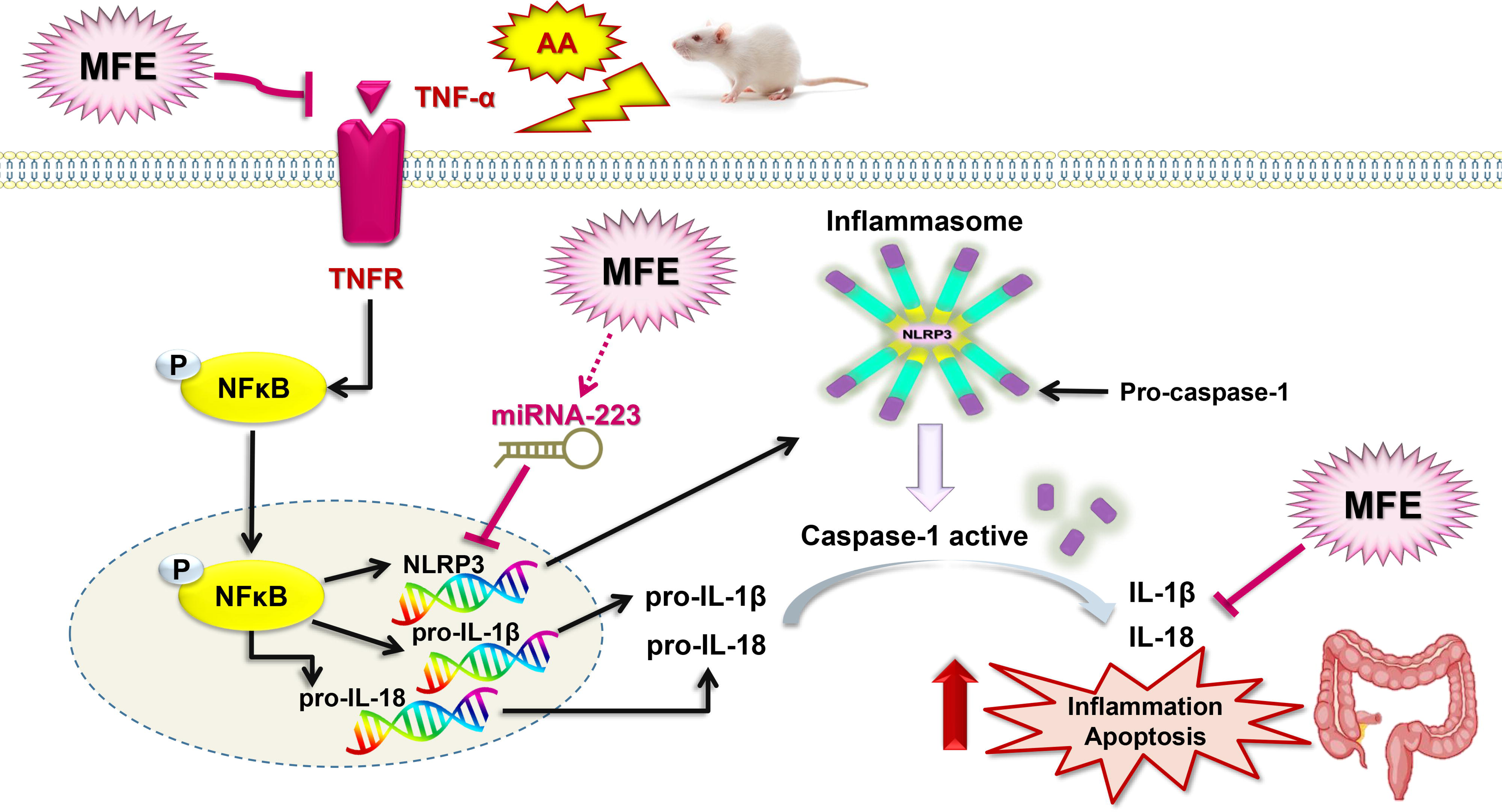

## Abbreviations

AA: acetic acid
DAI: disease activity index
GAE: gallic acid equivalents
H&E: Hematoxylin and Eosin
IBD: inflammatory bowel disease
IL: interleukin
MFE: mulberry fruit extract
NFκB: nuclear factor kappa B
NLRP3: nod-like receptor pyrin domain-1 containing 3
Sulfa: sulfasalazine
TAC: total anthocyanin content
TFC: total flavonoid content
TNFα: tumor necrosis factor alpha
TNFR: tumor necrosis factor receptor
TPC: total phenolic content
UA: ulcer area
UC: ulcerative colitis
UI: ulcer index

## Conflict of interest

The authors declare that they have no conflict of interest.

## Funding

This research did not receive any specific grant from funding agencies in the public, commercial, or not-for-profit sectors.

## Notes

### Competing Interest Statement

The authors have declared no competing interest.

